# Radiosynthesis, *in vitro* and preliminary *in vivo* evaluation of the novel glutamine derived PET tracers [^18^F]fluorophenylglutamine and [^18^F]fluorobiphenylglutamine

**DOI:** 10.1101/2020.02.14.948372

**Authors:** Tristan Baguet, Jeroen Verhoeven, Glenn Pauwelyn, Jiyun Hu, Patricia Lambe, Stef De Lombaerde, Sarah Piron, Sam Donche, Benedicte Descamps, Ingeborg Goethals, Christian Vanhove, Filip De Vos, M. Hassan Beyzavi

**Author notes:** **Corresponding Authors**, Tristan Baguet, Ottergemsesteenweg 460, 9000 Gent, Belgium, M. Hassan Beyzavi, Department of Chemistry and Biochemistry, Fayetteville, AR 72701, USA.

## Abstract

**Introduction:** Glucose has been deemed the driving force of tumor growth for decades. However, research has shown that several tumors metabolically shift towards glutaminolysis. The development of radiolabeled glutamine derivatives could be a useful molecular imaging tool for visualizing these tumors. We elaborated on the glutamine-derived PET tracers by developing two novel probes, namely [^18^F]fluorophenylglutamine and [^18^F]fluorobiphenylglutamine

**Materials and methods:** Both tracers were labelled with fluorine-18 using our recently reported ruthenium-based direct aromatic fluorination method. Their affinity was evaluated with a [^3^H]glutamine inhibition experiment in a human PC-3 and a rat F98 cell line. The imaging potential of [^18^F]fluorophenylglutamine and [^18^F]fluorobiphenylglutamine was tested using a mouse PC-3 and a rat F98 tumor model.

**Results:** The radiosynthesis of both tracers was successful with overall non-decay corrected yields of 18.46 ± 4.18 % (n=10) ([^18^F]fluorophenylglutamine) and 8.05 ± 3.25 % (n=5) ([^18^F]fluorobiphenylglutamine). *In vitro* inhibition experiments showed a moderate and low affinity of fluorophenylglutamine and fluorobiphenylglutamine, respectively, towards the human ASCT-2 transporter. Both compounds had a low affinity towards the rat ASCT-2 transporter. These results were endorsed by the *in vivo* experiments with low uptake of both tracers in the F98 rat xenograft, low uptake of [^18^F]FBPG in the mice PC-3 xenograft and a moderate uptake of [^18^F]FPG in the PC-3 tumors.

**Conclusion:** We investigated the imaging potential of two novel PET radiotracers [^18^F]FPG and [^18^F]FBPG. [^18^F]FPG is the first example of a glutamine radiotracer derivatized with a phenyl group which enables the exploration of further derivatization of the phenyl group to increase the affinity and imaging qualities. We hypothesize that increasing the affinity of [^18^F]FPG by optimizing the substituents of the arene ring can result in a high-quality glutamine-based PET radiotracer.

**Advances in Knowledge and Implications for patient care:** We hereby report novel glutamine-based PET-tracers. These tracers are tagged on the arene group with fluorine-18, hereby preventing *in vivo* defluorination, which can occur with alkyl labelled tracers (e.g. (2*S*,4*R*)4-[^18^F]fluoroglutamine). [^18^F]FPG shows clear tumor uptake *in vivo*, has no *in vivo* defluorination and has a straightforward production. We believe this tracer is a good starting point for the development of a high-quality tracer which is useful for the clinical visualization of the glutamine transport.

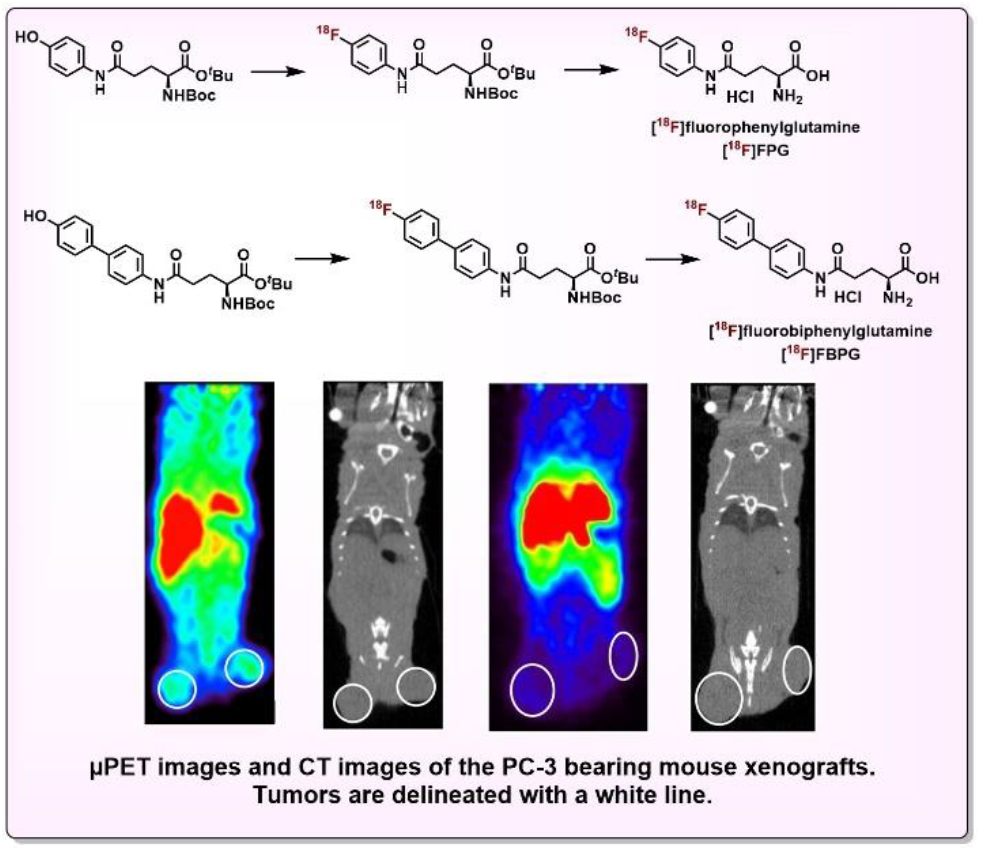

## 1. Introduction

The introduction of nuclear medicine in clinical practice has shifted the use of imaging in oncology. While magnetic resonance imaging (MRI) and computed tomography (CT) visualize tumor anatomy, the addition of nuclear medicine provides a more functional understanding. Positron emission tomography (PET) is a nuclear medicine technique for biomedical imaging that uses radiotracers interacting with some of the hallmarks in cancer, making it possible to visualize what is happening on a molecular level^1–3^. A successful example of such a PET radiotracer is [^18^F]fluorodeoxyglucose ([^18^F]FDG). Ever since its Federal Drug Agency (FDA) approval in 1996, [^18^F]FDG has been the golden standard in nuclear medicine imaging in oncology. Due to the high sensitivity, the application of [^18^F]FDG PET has proven to be useful, for example, in staging and therapy response assessment^2–4^. Nevertheless, [^18^F]FDG is of limited value for tumors with high background uptake (e.g. brain tumors), low tumor uptake (e.g. neuroendocrine tumors, prostate cancer, hepatocellular carcinoma) or in the presence of tumor-associated inflammation^5–8^.

Amino acids also show increased uptake in malignant cells and seem to overcome several of the glucose-related limitations. Several radiolabeled amino acids targeting the L-type amino acid transporter 1 (LAT1) such as O-(2-^18^F-fluoroethyl)-L-tyrosine ([^18^F]FET), (*S*-^11^C-methyl)-L-methionine ([^11^C]MET) and 3,4-dihydroxy-6-^18^F-fluoro-L-phenylalanine ([^18^F]FDOPA) show favorable tumor visualization and are already used in clinical practice^5–8^. Another amino acid that draws attention in cancer metabolism is glutamine^9^. Being the most abundant amino acid in blood and muscle, glutamine is also an essential amino acid for malignant cells as several tumors seem to be unable to survive without the presence of exogenous glutamine. It is both a carbon and nitrogen source and participates in the cellular redox homeostasis, which explains the “glutamine addiction” ^10–13^. Some tumors have even been reported to metabolically shift to glutaminolysis, resulting in negative [^18^F]FDG PET scans^14,15^. All these aspects led to the development of glutamine-based PET tracers e.g. [^11^C]glutamine, (2*S*,4*R*)4-[^18^F]fluoroglutamine, (2*S*,4*S*)4-(3-[^18^F]fluoropropyl)glutamine and (2S,4S)2,5-diamino-4-(4-(2-[^18^F]fluoroethoxy)benzyl)-5-oxopentanoic acid ^16–20^. As an isotopic variant, [^11^C]glutamine has identical physicochemical properties to normal glutamine. However, due to its short half-life (t_1/2_ = 20.3 min) and difficult radiochemical synthesis, no human clinical studies have yet been performed. To address the short half-life, a fluorine-18 analogue was developed: (2*S*,4*R*)4-[^18^F]fluoroglutamine. Unfortunately, the fluorine-18 derivative of glutamine shows *in vivo* defluorination and its synthesis is complex with different stereoisomers being formed. (2*S*,4*S*)4-(3-[^18^F]-fluoropropyl)glutamine is more stable *in vivo*, but suffers low labelling yields and shows system L preference. (2*S*,4*S*)2,5-diamino-4-(4-(2-[^18^F]-fluoroethoxy)benzyl)-5-oxopentanoic acid shows much lower uptake *in vitro* and was not yet evaluated *in vivo*^17–20^.

In this study, we further elaborated on the glutamine-derived PET-tracers. Therefore, we aimed to obtain fluorine-18 labelled inhibitors of the alanine-serine-cysteine transporter-2 (ASCT-2), the most important glutamine transporter. The first radiotracer we synthesized was the fluorine-18 variant of the known inhibitor L-γ-glutamyl-p-nitroanilide (GPNA)^21^. For the second radiotracer we decided to expand the aromatic system of GPNA to a biphenyl group. As such, we aimed at a better affinity for the ASCT-2 transporter by better filling the hydrophilic pocket described in different homology models^21–23^. The organic synthesis of both precursors and cold reference products was based on a method described by Schulte et al.^22^. In order to avoid interference with the glutamine backbone, implementation of the radioisotope was performed directly at the aromatic rings. Therefore, we attempted to develop the fluorine-18 labeled derivative of glutamine via a ruthenium-based direct aromatic fluorination method. Both tracers were evaluated *in vitro* for their affinity towards the glutamine transporters. Finally, as a proof of concept, PET-images were acquired in a mouse model for prostate cancer and a rat model for glioblastoma.

## 2. Materials & methods

### a. General

All reactions described were performed under nitrogen atmosphere. Chemicals were at least reagent grade, obtained from Sigma Aldrich (Bornem, Belgium), TCI Europe (Zwijndrecht, Belgium), Activate Scientific (Prien, Germany), Acros (Geel, Belgium) or ChemCollect (Wuppertal, Germany) and used as received. All solvents were obtained from Chemlab (Zedelgem, Belgium) and were at least HPLC-grade. Pre-coated TLC sheets ALUGRAM^®^ Xtra SIL G/UV_254_ were used to monitor reactions under 254 nm UV-light. Staining of TLC was performed by spraying with either basic aqueous solution of KMnO_4_ (1.0 g KMnO_4_, 2.0 g K_2_CO_3_, 100 mL H_2_O) or ninhydrin (1.5g ninhydrin, 3.0 mL acetic acid, 100 mL EtOH) solution and subsequent charring. Purification was performed with silica column chromatography either manually with Machery-Nagel Kieselgel 60 (63-200 μm) or on a Reveleris X2 (Grace) automatic system and accessory pre-packed silica columns. A Waters LCT Premier XE Time of Flight (TOF) mass spectrometer equipped with electrospray ionization (ESI) and a modular Lockspray^TM^ interface was used to obtain high resolution mass spectra. NMR spectra were recorded with a Varian Mercury-300BB (300/75 MHz) spectrometer.

### b. Chemistry

#### General procedure A (amine acylation of aniline with protected glutamate)

To a heat-dried flask containing the Boc-l-Glu-OtBu (500 mg, 1.65 mmol, 1.00 eq), aniline (1.65 mmol, 1.00 eq), 1-[Bis(dimethylamino)methylene]-1H-1,2,3-triazolo[4,5-b]pyridinium 3-oxid hexafluorophosphate (HATU) (627 mg, 1.65 mmol, 1.00 eq), diisopropylethylamine (DIPEA) (0.57 mL, 3.30 mmol, 2.00 eq) and a magnetic stirring bar under N_2_ gas was added anhydrous DMF (18 mL, 11mL/ mmol) as a solvent. The reaction mixture was stirred overnight at 120 °C. The next day the solution was cooled to room temperature and DMF was evaporated *in vacuo*. Next, the mixture was partitioned in water/ CH_2_Cl_2_, and extracted with CH_2_Cl_2_ (2x). The organic layer was dried by passing it over a Büchner funnel filled with sodium sulphate. The filtrate was absorbed onto celite^®^, concentrated *in vacuo* and purified by flash chromatography on silica gel (hexane:ethyl acetate).

#### General procedure B (cleavage of tert-butyl and N-boc protection groups)

The purified compounds were dissolved in 4.0M HCl in dioxane (10 mL/ mmol) in an anhydrous environment and were stirred 4 hours at 40 °C. The reaction mixture was concentrated *in vacuo* and used without further purification.

#### *tert*-Butyl N2-(*tert*-butoxycarbonyl)-N5-(4-fluorophenyl)-L-glutaminate (1)

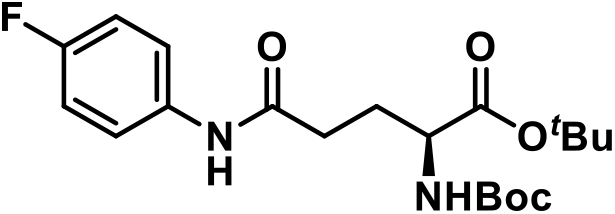

Boc-l-Glu-OtBu (500 mg, 1.65 mmol) and 4-fluoroaniline (183 mg, 1.65 mmol) were coupled using procedure A. The mixture, absorbed onto celite^®^, was purified using automated flash chromatography (12→ 25% EtOAc in hexane) to afford **1** (503 mg, 1.27 mmol) as a white powder in 77% yield. R_f_ 0.35 (25% EtOAc/75% Hex). ^1^H NMR (300 MHz, CDCl_3_): δ 1.45 (d, *J*=3.0 Hz, 18H, *t*Bu), 1.74-1.94 (m, 1H, CH_2_-<u>CH</u>_2_-CH), 2.18-2.32 (m, 1H, CH_2_-<u>CH</u>_2_-CH), 2.36-2.48 (m, 2H, NH-OC-CH_2_-<u>CH</u>_2_), 4.14-4.28 (m, 1H, <u>CH</u>-NH), 5.36 (d, *J*=7.7 Hz,1H, NH-*t*Bu), 6.95-7.04 (m, 2H, H_Phe_), 7.57 (dd, *J*=4.8 Hz and 3.8 Hz, 2H, H_Phe_), 8.93 (s, 1H, PhNH). ^13^C NMR (75 MHz, CDCl_3_): δ 27.96, 28.30, 30.69, 34.02, 53.19, 80.55, 82.79, 115.29, 115.58, 121.31, 134.54, 156.65, 157.52, 160.74, 170.48, 171.22. ^19^F NMR (282 MHz, DMSO-*d*6): δ [-119.69; −119.56] (m). Exact Mass: (ESI-MS) for C_20_H_30_FN_2_O_5_ [M+H] found, 397.2161; calcd, 397.2139.

#### tert-Butyl N2-(tert-butoxycarbonyl)-N5-(4-hydroxyphenyl)glutaminate (2)

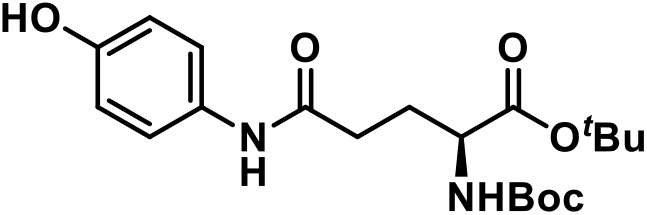

Boc-l-Glu-OtBu (500 mg, 1.65 mmol) and 4-aminophenol (180 mg, 1.65 mmol) were coupled using procedure A. The mixture, absorbed onto celite^®^, was purified using automated flash chromatography(25→ 50% EtOAc in hexane) to afford **2** (382 mg, 0.97 mmol) as a red crystalline powder in 59% yield. Rf 0.46 (50% EtOAc/ 50% Hex). ^1^H NMR (300 MHz, CDCl_3_): δ 1.44(s, 18H, *t*Bu), 1.81-1.98 (m, 1H, CH_2_-<u>CH</u>_2_-CH), 2.15-2.29 (m, 1H, CH_2_-<u>CH</u>_2_-CH), 2.40 (t, *J*=7.0 Hz, 2H, O=C-<u>CH</u>_2_-CH_2_), 2.81(s, 1H, OH_Phe_), 4.07-4.25 (m, 1H, <u>CH</u>-NH), 5.38 (d, *J*=8.0 Hz, 1H, NH-*t*Bu), 6.78(d, *J*=8.7Hz, 2H, H_Phe_), 7.36(d, *J*=8.5Hz, 2H, H_Phe_), 8.55(s,1H, PhNH). ^13^C NMR (75 MHz, CDCl_3_): δ 27.95, 28.30, 30.21, 33.81, 53.41, 80.50, 82.72, 115.63, 122.16, 130.58, 153.26, 156.49, 170.75, 171.33. Exact Mass: (ESI-MS) for C_20_H_31_N_2_O_6_ [M+H] found, 395.2174; calcd, 395.2182.

#### tert-Butyl N2-(tert-butoxycarbonyl)-N5-(4’-fluoro-[1,1’-biphenyl]-4-yl)glutaminate (3)

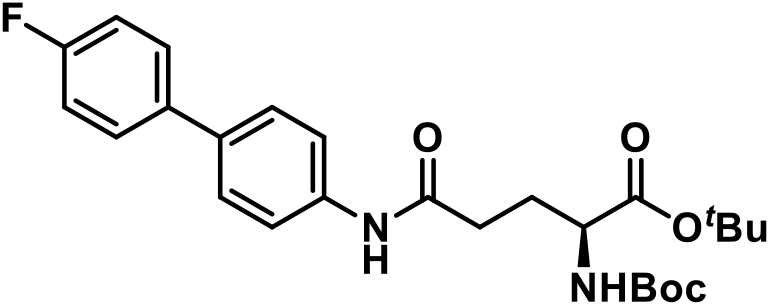

Boc-l-Glu-OtBu (500 mg, 1.65 mmol) and 4’-fluoro-[1,1’-biphenyl]-4-amine (309 mg, 1.65 mmol) were coupled using procedure A. The mixture, absorbed onto celite^®^, was purified using manual flash chromatography (15→ 30% EtOAc in hexane) to afford **3** (577 mg, 1.22 mmol) as a white powder in 74% yield. Rf 0.19 (25% EtOAc/ 75% Hex). ^1^H NMR (300 MHz, CDCl_3_): δ 1.47 (d, *J*=6.0 Hz, 18H, *t*Bu), 1.79-1.95 (m, 1H, CH_2_-<u>CH</u>_2_-CH), 2.21-2.36 (m, 1H, CH_2_-<u>CH</u>_2_-CH), 2.46(t, *J*=7.3 Hz, 2H, O=C-<u>CH</u>_2_-CH_2_), 4.174.30 (m, 1H, <u>CH</u>-NH), 5.37(d, *J*=8.0Hz, 1H, NH-*t*Bu), 7.06-7.15 (m, 2H, H_Phe_), 7.47-7.56 (m, 4H, H_Phe_), 7.69 (d, *J*=8.4Hz, 2H, H_Phe_), 8.96 (s,1H, PhNH). ^13^C NMR (75 MHz, CDCl_3_): δ 27.98, 28.31, 30.80, 34.19, 53.21, 80.56, 82.80, 115.43, 115.72, 120.06, 127.38, 128.27, 128.37, 135.84, 136.79, 137.77, 156.65, 160.64, 163.91, 170.52, 171.25. ^19^F NMR (300 MHz, CDCl_3_): δ: −116.22 (s). Exact Mass: (ESI-MS) for C_26_H_34_FN_2_O [M+H] found, 473.2456; calcd, 473.2452.

#### tert-Butyl N2-(tert-butoxycarbonyl)-N5-(4’-hydroxy-[1,1’-biphenyl]-4-yl)glutaminate (4)

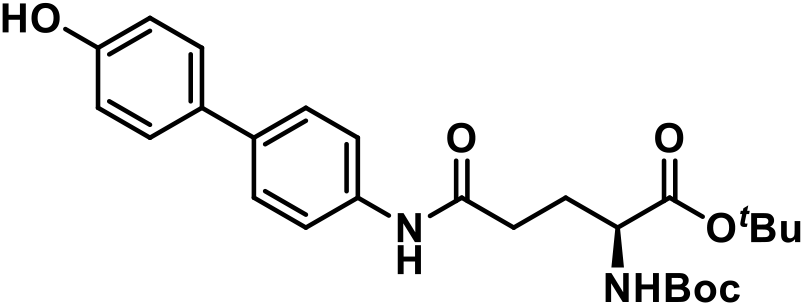

Boc-l-Glu-OtBu (500 mg, 1.65 mmol) and 4’-amino-[1,1’-biphenyl]-4-ol (305 mg, 1.65 mmol) were coupled using procedure A. The mixture, absorbed onto celite^®^, was purified using automated flash chromatography (20→ 50% EtOAc in hexane) to afford **4** (442 mg, 0.94 mmol) as a white powder in 57% yield. Rf 0.37 (50% EtOAc/ 50% Hex). ^1^H NMR (300 MHz, DMSO-*d*6): δ 1.38 (d, *J*=5.7 Hz, 18H, *t*Bu), 1.72-1.88 (m, 1H, <u>CH</u>_2_-CH_2_-CH), 1.92-2.07 (m, 1H, CH_2_-<u>CH</u>_2_-CH), 2.38 (t, *J*=7.5 Hz, 2H, O=C-<u>CH</u>_2_-CH_2_), 3.72-3.89 (m, 1H, <u>CH</u>-NH), 6.70-6.84 (m, 2H, H_Phe_), 7.13 (d, *J*= 7.8 Hz, 1H, H_Phe_), 7.39-7.51 (m, 3H, H_Phe_), 7.57-7.63 (m, 2H, H_Phe_), 9.45 (s, 1H, PhNH), 9.94 (s, 1H, PhOH). ^13^C NMR (75 MHz, DMSO-*d*6) δ 26.68, 28.10, 28.63, 33.12, 54.36, 78.52, 80.78, 116.10, 119.80, 126.48, 127.16, 127.70, 131.02, 135.30, 138.21, 155.99, 157.18, 170.67, 172.08. Exact Mass: (ESI-MS) for C_26_H_35_N_2_O_6_ [M+H] found, 471.2487; calcd, 471.2495

#### N5-(4-fluorophenyl)glutamine (5)

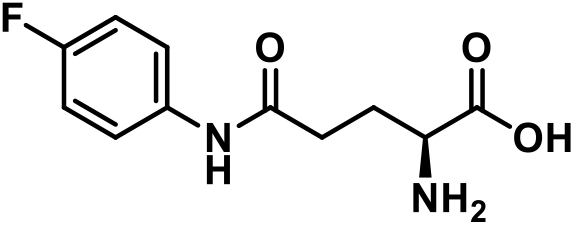

The *tert*-butyl and *N*-boc protection groups of compound 1 were removed using procedure B, to obtain compound **5**, a white powder, in quantitative yields. ^1^H NMR (300 MHz, DMSO-*d*_6_): δ 2.00-2.15 (m, 2H, CH_2_-<u>CH</u>_2_-CH), 2.51-2.65 (m, 2H, O=C-<u>CH</u>_2_-CH_2_), 3.80 (t, *J*=6.2Hz, 1H, <u>CH</u>-NH2), 7.10 (t, *J*=8.9 Hz, 2H, H_Phe_), 7.61 (dd, *J=* 9.0 Hz and 5.1 Hz, 2H, H_Phe_), 10.35 (s, 1H, PhNH). ^13^C NMR (75 MHz, DMSO-*d_6_*): δ 26.39, 32.16, 52.10, 115.45, 115.74, 121.15 (m), 136.03 (m), 156.71, 159.88, 170.25, 170.38, 171.18. ^19^F NMR (282 MHz, DMSO-D6): δ [-119.70; −119.56] (m). Exact Mass: (ESI-MS) for C_11_H_14_FN_2_O_3_ [M+H] found, 241.0987; calcd, 241.0988.

#### N5-(4’-fluoro-[1,1’-biphenyl]-4-yl)glutamine (6)

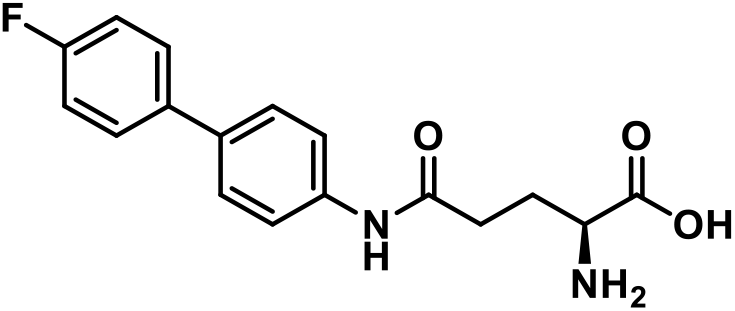

The *tert*-butyl and *N*-boc protection groups of compound **3** were removed using procedure B, to obtain compound **6**, a white powder, with quantitative yields. ^1^H NMR (300 MHz, TFA-*d*): δ 2.47-2.88 (m, 2H, CH_2_-<u>CH</u>_2_-CH), 3.07-3.23 (m, 2H, O=C-<u>CH</u>_2_-CH_2_), 4.58-4.76 (m, 1H, <u>CH</u>-NH2), 7.09-7.78(m, 8H, H_Phe_). ^13^C NMR (75 MHz, TFA-D): δ 24.77, 32.13, 53.59, 66.25, 114.87, 115.16, 123.78, 127.48, 128.33 (d, *J=*8Hz), 132.57, 135.99 (d, *J=*3Hz), 140.63, 164.31, 172.54, 174.70. ^19^F NMR (300 MHz, DMSO-*d_6_*): δ [-116.24; −116.10] (m). Exact Mass: (ESI-MS) for C_17_H_18_FN_2_O_3_ [M+H] found, 317.1297; calcd, 317.1301

### c. Radiochemical synthesis of [^18^F]fluorphenylglutamine ([^18^F]FPG) and [^18^F]fluorbiphenylglutamine ([^18^F]FBPG)

The chemical labelling of compounds **2** and **4** with fluorine-18 was done via an aromatic fluorination method described by Beyzavi et al.^24^. The fluoride used for these reactions was produced in a cyclone 18/9 cyclotron (IBA) with a ^18^O(p,n) nuclear reaction. The [^18^F^-^] (2.2 GBq) obtained in an aqueous solution was trapped on a HCO_3_^-^ SPE column and washed with 1.0 mL extra dry acetonitrile. A mixture of precursor (3.0mg), CpRu(COD)Cl (5.0 mg) and *N*,*N*’-bis(2,6-diisopropylphenyl)-2-chloroimidazolium chloride) (18.0 mg) was used to elute the fluoride from the SPE column in a 5.0 mL borosilicate vial. Extra dry DMSO (150 μL) and MeCN:DMSO (50:50, 50 μL) was used to flush the remaining fluoride. The resulting mixture was heated to 125°C for 30 minutes.

Upon completion of the reaction, the vial was cooled in an ice bath to room temperature. After cooling down the mixture was diluted with ultrapure water (4.5 mL) and trapped on a Sep-Pak C18 cartridge (preconditioned with MeCN (10.0 mL) and ultrapure water (10.0 mL)). The cartridge was washed with ultrapure water (10.0 mL), flushed twice with air and eluted with extra dry MeCN (1.0 mL). The MeCN was evaporated under a gentle stream of nitrogen at 125 °C. After evaporation of the solvent, the vial was cooled to room temperature. The protecting groups were removed by adding 4 M HCl in dioxane (200 μL) and heating the vial for 3.5 minutes to 90°C. After cooling down the mixture, mobile phase (300 μL) was added to neutralize the pH. Purification was then done by injecting onto HPLC using a C18 prep column (Atlantis T3 OBD prep column, 100 Å, 10 μM, 10mm × 250mm, 1/pkg, Waters). Elution of the desired radiotracer was followed with a radiodetector (Ludlum Measurements Inc) coupled to the HPLC. [^18^F]FPG eluted after approximately 12 minutes (mobile phase ethanol/ ammonium acetate (10mM) in ultrapure water 10:90 (V:V); flowrate 6 mL/min) and was used as such. [^18^F]FBPG eluted after approximately 16 minutes (mobile phase ethanol/ ammonium acetate (10 mM) in ultrapure water 30:70 (V:V); flowrate 6 mL/min) and was diluted with PBS before further usage.

Quality control of the obtained radiotracer was done by means of analytical HPLC with radio-and UV detection (205 nm; Waters) using a 1.0 mL/min flow and a Grace Alltima C18 (4.6×250 mm; 5 μm) column. The retention times of the fluorine-18 labelled compounds were compared to the retention time of the fluorine-19 reference compounds. [^18^F]FPG was compared to [^19^F]FPG (retention time 8.53 minutes) in an isocratic run using ethanol/ ammonium acetate (10 mM) in ultrapure water 10:90 (V:V). [^18^F]FBpG was compared to [^19^F]FBPG (retention time 10.35 minutes) in an isocratic run using ethanol/ ammonium acetate (10 mM) in ultrapure water 30:70 (V:V). The LogD_7.4_ of both tracers were determined by adding 1 MBq of respectively [^18^F]FPG and [^18^F]FBPG to a mixture of PBS water and n-octanol (1:1; V:V). Samples were vortexed and centrifuged (10 min; 1100 g) where after aliquots were taken from each layer which were measured in a NaI (Tl) scintillation counter (Capintec; Ramsey, NJ, USA). LogD7.4 was calculated as the logarithm of the proportion of counts in the octanol and PBS layers.

### d. In vitro experiments

#### General information

The PC-3 and F98 cell lines were purchased from ATCC and the SLC1A5 knock-out cell line from Creative Biogene. The PC-3 cell line was cultivated in RPMI medium enriched with 10% fetal calf serum, 1% l-glutamine and penicillin/streptomycin (50 U/mL). Both the F98 cell line and SLC1A5 knock-out cell line were cultivated in Dulbecco’s modified Eagle’s medium (DMEM) enriched with 10% fetal calf serum, 1% l-glutamine and penicillin/streptomycin (50 U/mL). All cell lines were stored in an incubator set to 37 °C and 5% CO_2_ environment. The presence of ASCT-2 expression in PC-3 and F98 cell lines was verified with flow cytometry. The analysis was performed using the LSR II (BD Biosciences) as described by De Munter et al.^35^. Both cell lines (100,000 cells/ 100 μL) were stained at their surface with a primary (rabbit ab; ASCT-2 (V501)) and secondary antibody (Fab2 (PE Conjugate); anti-rabbit IgG) against ASCT-2 as a negative control. The cells were fixed and permeabilized followed by intracellular binding with the primary antibody. After binding of the primary antibody, the secondary antibody was added and after the incubation period, all samples were analyzed. The ASCT-2 knock-out cell line was engineered with CRISPR-CAS 9 using a HEK293 cell line. Absence of ASCT-2 was also explored via flow cytometry. The percentage of ASCT-2 positive cells was < 1%, indicating that the knock-out efficiency approximates 100%.

Cell tests were performed in 24-well plates (VWR, US). PC-3, F98 and SLC1A5 KO cells were seeded the day prior to the test in a concentration of respectively 175,000; 125,000 and 175,000 cells per milliliter. The 24-well plates used for the cell tests with the knock-out cell line were coated prior to seeding to achieve good attachment of the cells. To coat the plate 300 μL of Poly-D-lysine stock solution (0.1 mg/mL) was added to each well. The plates were dried for 30 minutes until all solvent was removed. Upon removal of the solvents the plates were ready for cell attachment. The [^3^H]glutamine uptake studies were performed in HEpES+ buffer (pH 7.4; 100 mM NaCl (Sigma Aldrich, Belgium), 2 mM KCl (Sigma Aldrich, Belgium), 1 mM MgCl_2_ (VWR, US), 1 mM CaCl_2_ (VWR, US), 10 mM Hepes (Sigma Aldrich, Belgium), 5 mM Tris (VWR, US), 1 g/L glucose (VWR, US) and 1 g/L Bovine Serum Albumin (Sigma Aldrich, Belgium)). A glutamine stock concentration (43.8 mg/ 30mL HEPES+) was used to make the desired concentration series. To each stock solution a total of 111 kBq [^3^H]-l-glutamine (Perkin Elmer, Massachusetts)/mL was added. Concentration series containing the compounds of interest ([^19^F]FPG and [^19^]FBPG) were made next to the reference glutamine concentration series. At the start of the experiments, the medium was aspirated, and all wells were washed twice with HEPES+. After removal of HEPES+, 250 μL test solution was added, and the plates were incubated for 5 minutes at 37°C. Uptake was stopped after 5 minutes by placing the plates on ice and adding 1 mL ice-cold PBS +BSA (1g/100mL). PBS +BSA was removed and the wells were washed twice with 2 mL ice cold PBS. Cells were lysed by placing the plates on a shaker after adding 250 μL 0.1 M NaOH (VWR, US). Aliquots of 150 μL were pipetted out of each well into a scintillation bottle (Perkin Elmer, Massachusetts, USA). To each scintillation bottle, 5 mL of scintillation cocktail (Ultima Gold, Perking Elmer, Massachusetts, USA) was added. Samples were measured with an automated scintillation counter (TriCarb 2900 TR; Perkin Elmer, Massachusetts, USA). From each plate 25 μL cell lysis was subjected to the bichoninic acid assay (BCA) (ThermoFisher Scientific) to correct for protein density.

##### Concentration dependency

The uptake of [^3^H]-l-glutamine was tested in F98 rat cells and PC-3 human cells in absence of cold reference product and was compared to the uptake in the presence of cold reference products. The glutamine stock solution used ranged from 10 μM to 1200 μM. To data obtained from the liquid scintillation counter was plotted via non-linear regression in GraphPad prism v5.01 (GraphPad Software, San Diego, CA, USA) to obtain Michealis-Menten parameters. K_m_ (values in absence of cold product) and K_m,app_ (apparent values in presence of cold product) data were used to calculate the Ki values according to the formula: *Ki* = [*I*]/((*Km_app_*/*Km*) –1) with [I] being the concentration of inhibitor.

##### Inhibition [^3^H]-l-glutamine uptake in a knock-out cell line

The inhibition of the [^3^H]-l-glutamine uptake in absence of ASCT-2 was measured in a SLC1A5 knock-out cell line to evaluate the selectivity of both radiotracers for ASCT-2. The cells were incubated with 250 μL 50 μM [^3^H]-l-glutamine/glutamine in the absence and presence of both tracers for 5 minutes at 37°C.

#### In vitro evaluation of stability in formulation

The stability of [^18^F]FPG and [^18^F]FBPG was evaluated in their formulation. After radiosynthesis, the radiotracer was incubated in its formulation for 30, 70, 120 and 180 minutes at 37 °C. At the indicated time points 7 MBq was analyzed by the same analytical HPLC method used to determine the purity and identity of [^18^F]FPG and [^18^F]FBPG.

### e. In vivo experiments

All animal experiments were in accordance with the regulations set up by the Ghent University Ethical Committee and were approved prior to the start of the study by the local Ethical Committee on Animal Experiments (ECD 17/08 and ECD 18/21). During housing water and food was provided *ad libitum*. Animals were fasted minimal six hours prior to PET scans. Anesthesia was induced with 5.0% isoflurane (V:V in O_2_) and maintained with 2.0% until the end of the experiment. After intravenous injection in the lateral tail vein, 1-hour dynamic PET acquisitions were performed on a FLEX Triumph II small animal PET/CT-scanner (PET field of view: 7.5 cm axial; 1.3 mm spatial resolution; TriFoil Imaging). Upon completion of the PET-scan an additional CT-scan was performed for anatomical correlation purposes. [^18^F]FPG and [^18^F]FBPG uptake was evaluated in both PC-3 and F98 tumors. The PET data were obtained in list mode and reconstructed iteratively (50 iterations) in frames of one minute the first twenty-five minutes and frames of five minutes for the following thirty-five minutes. Region of interests (ROI) were drawn manually around the tumor and tissue for comparison (leg muscle and contralateral brain tissue for the PC-3 and F98 animal models, respectively). ROI’s were converted to standard uptake values (SUV) following

> *SUV* = (*Radioactivity in ROI/ injected activity*) × *body weight*

with injected activity corrected for radioactive decay and residual radioactivity in the syringe.

#### PC-3 xenograft model

At the day of inoculation, the PC-3 cells were harvested and suspended in non-enriched RPMI medium to obtain a 5 × 10^7^ cells/ mL cell suspension. Male swiss nu/nu mice (n=5) were inoculated with 5 × 10^6^ pC-3 tumor cells (+0,1 mL Matrigel^®^) in the right flank at the age of 8 weeks. One week later the mice were inoculated in the left flank with 5 × 10^6^ PC-3 tumor cells (+0,1 mL Matrigel^®^). Tumor growth was followed with manual caliper measurement. Dynamic μPET scans were acquired seven weeks post-inoculation after intravenous injection of 18.5 MBq radiotracer. PET and CT images were aligned using the AMIDE (Source Forge Media, La Jolla, CA, USA) software. Tumor-to-muscle (T/M) ratios were calculated with the leg muscle as reference region.

#### Orthotopic F98 model

The F98 glioblastoma rat model was obtained by the same method described by Verhoeven et al.^36^. A suspension of 20 × 10^3^ F98 tumor cells was injected into the right frontal lobe of female Fischer rats (n=3). Tumor growth was visualized using T2-weighted and contrast-enhanced T1-weighted MRI (7T pharmaScan 70/16, Bruker BioSpin, Ettlingen, Germany). Upon tumor confirmation on MRI (± 12 days post inoculation), dynamic PET acquisitions were obtained after injection of 37 MBq radiotracer. Coregistration of the images was done in PMOD. First, the CT-MRI and CT-PET images were aligned. Next the MRI images were aligned to the PET images. Tumor tissue was delineated based on the contrast-enhancing region on T1-weighted MRI. A reference region (2 × 2 × 2 mm^3^) was delineated contralateral to the tumor.

## 3. Results

### a. Chemistry

The synthetic approach for both the precursors and cold reference products of [^18^F]fluorophenylglutamine and [^18^F]fluorobiphenylglutamine is based on the method described by Schulte et al. (2015) (scheme 1)^22^. Formation of the amide was realized by a HATU based coupling of the aniline to the protected glutamate. Subsequent deprotection of 1 and 3 with 4M HCl in dioxane provides the cold reference compounds 5 and 6. Deprotection of 1 and 3 was tested with aqueous HCl (6 M), however, this led to excessive generation of impurities. The use of 4.0 M HCl in dioxane led to pure cold reference products.

**Scheme 1:**
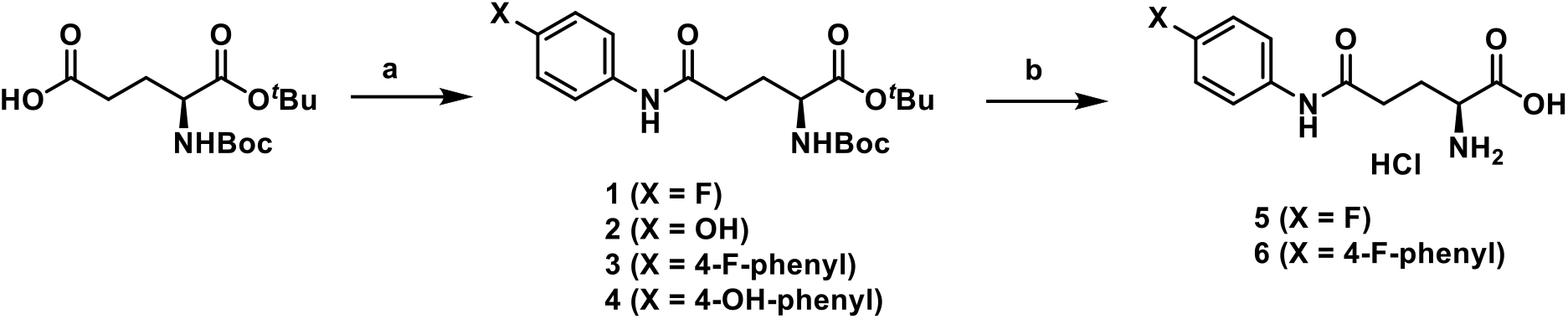
Synthesis of precursors for radiolabeling (2 and 4) and cold reference products (5 and 6). Reagents and conditions: a) aniline, HATU, DIPEA, DMF, 120°C, overnight; b) HCl (4M/dioxane), 40°C, 4h.

### b. Radiochemistry

#### [^18^F]fluorophenylglutamine

[^18^F]fluorophenylglutamine ([^18^F]FPG) was synthesized as described in scheme 2 with a non-decay corrected radiochemical yield of 18.46 ± 4.18 % (n=10) and a total synthesis time of 88 ± 4 minutes. Compound verification was performed by means of co-injecting cold reference product *N5-(4-fluorophenyl)glutamine* onto analytical HPLC (t_r_ [^18^F]FPG = 8.8 minutes) (chromatogram see supplementary information Figures S1 and S2). Radiochemical purity of [^18^F]FPG was >99% as determined by analytical HPLC. The logD_7.4_ (octanol/pBS ratio at pH 7.4) measured for [^18^F]FPG was −0.70 ± 0.02 (n=3).

#### [^18^F]fluorobiphenylglutamine

[^18^F]fluorobiphenylglutamine ([^18^F]FBPG) was synthesized based on our recent protocol^24^ as described in scheme 2 with a non-decay corrected radiochemical yield of 8.05 ± 3.25 % (n=5) and a total synthesis time of 97 ± 12 minutes. Compound verification was done by means of co-injecting cold reference product *N5*-(*4’-fluoro-[1,1’-biphenyl]-4-yl*)*glutamine* onto analytical HPLC (t_r_ [^18^F]FBPG = 11.1 minutes) (chromatogram see supplementary information Figure S3 and S4). Radiochemical purity of [^18^F]fluorobiphenylglutamine was >99% as determined by analytical HPLC. The LogD7.4 (octanol/PBS ratio at pH 7.4) measured for [^18^F]FBPG was 1.11 ± 0.01 (n=3).

**Scheme 2:**
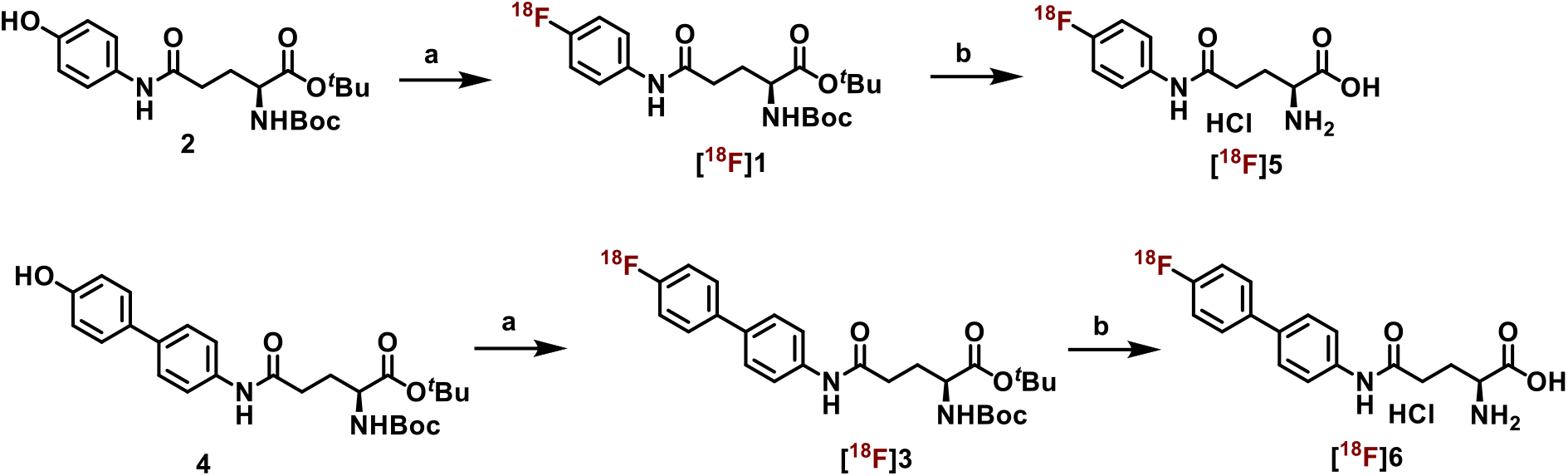
Radiochemical synthesis of [^18^F]FPG and [^18^F]FBPG. Reagents and conditions: a) Ruthenium complex: CpRu(COD)Cl (Cp: cyclopentadienyl, COD: 1,5-cyclooctadiene, N,N’-bis(2,6-diisopropylphenyl)-2-chloroimidazolium chloride, [^18^F]fluoride, DMSO:MeCN (1:1); b) HCl (4M/dioxane).

### c. In vitro experiments

#### Flow cytometry

Flow cytometry confirmed the expression of ASCT-2 on both the PC-3 and F98 cell line (see Figure 1). The PC-3 cell line showed a median population of 12,056 while the median population of the negative control was 576. In comparison, the median population in the F98 was 28,585 with a median population of the negative control of 919.

**Figure 1:**
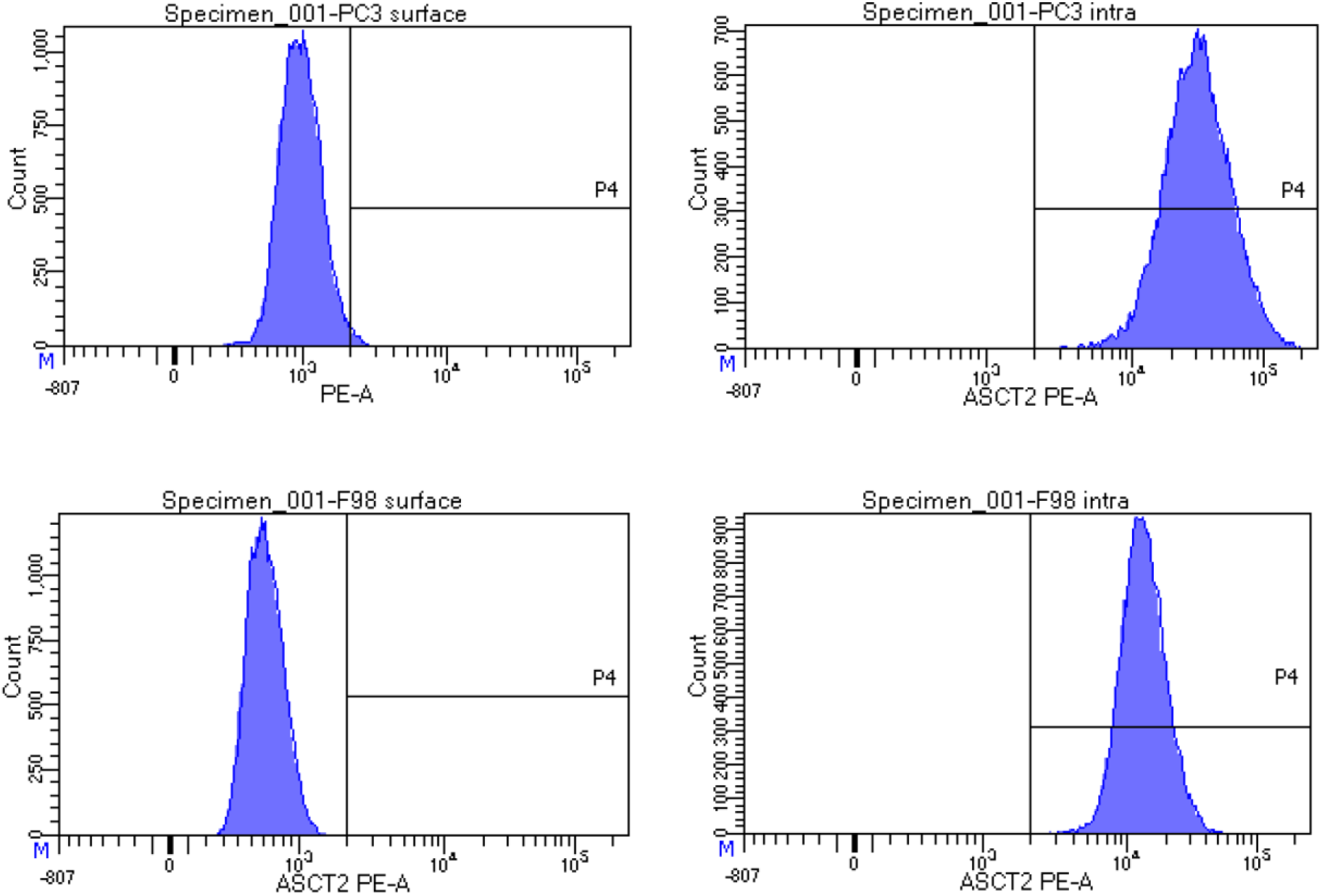
FACS analysis of ASCT-2 presence on PC-3 and F98 cells. The top left figure represents surface staining (negative control). The top right figure is intracellular staining of PC-3 cells. The bottom left figure is surface staining of F98 cells (negative control). The bottom right figure is intracellular staining of F98 cells.

#### Concentration dependency

Affinity constants Ki were calculated for both compounds and can be found in Table 1^1^. FPG shows moderate affinity towards the ASCT-2 transporter on the human PC-3 cell line and a low affinity to the transporter expressed on the rat F98 cell line. FBPG has low affinity to the transporter on both PC-3 and F98 cell lines. Michaelis-Menten plots for glutamine uptake in the presence and absence of the inhibitors can be found in Figure 2.

**Figure 2:**
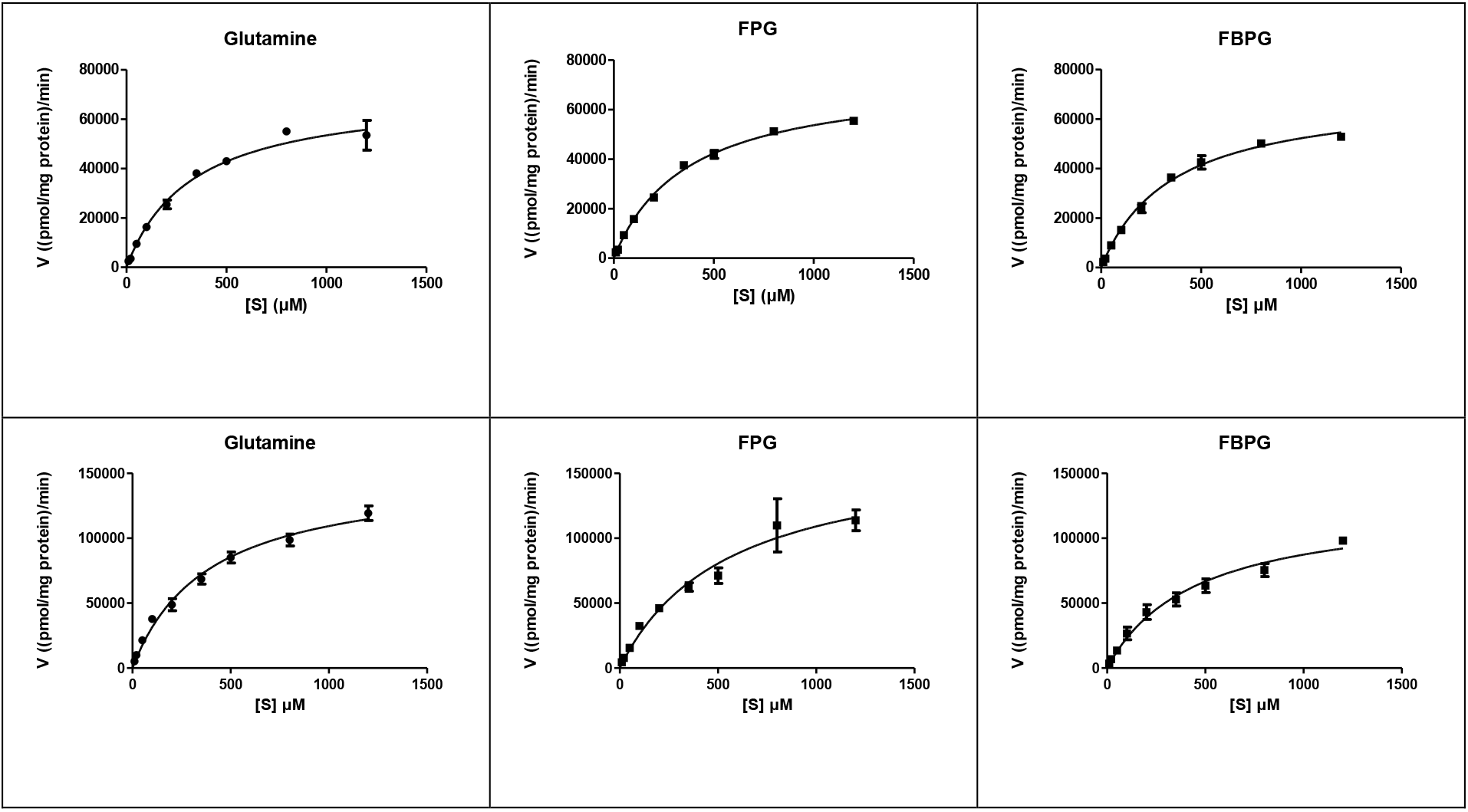
Concentration dependent uptake of [^3^H]glutamine in F98 cell line (top row) and PC-3 cell line (bottom row). Top left and bottom left graphs represent baseline uptake of [^3^H]glutamine. The middle graphs represent uptake in presence of FPG. Top and bottom right graphs show the uptake of [^3^H]glutamine in the presence of FBPG.

**Table 1:**
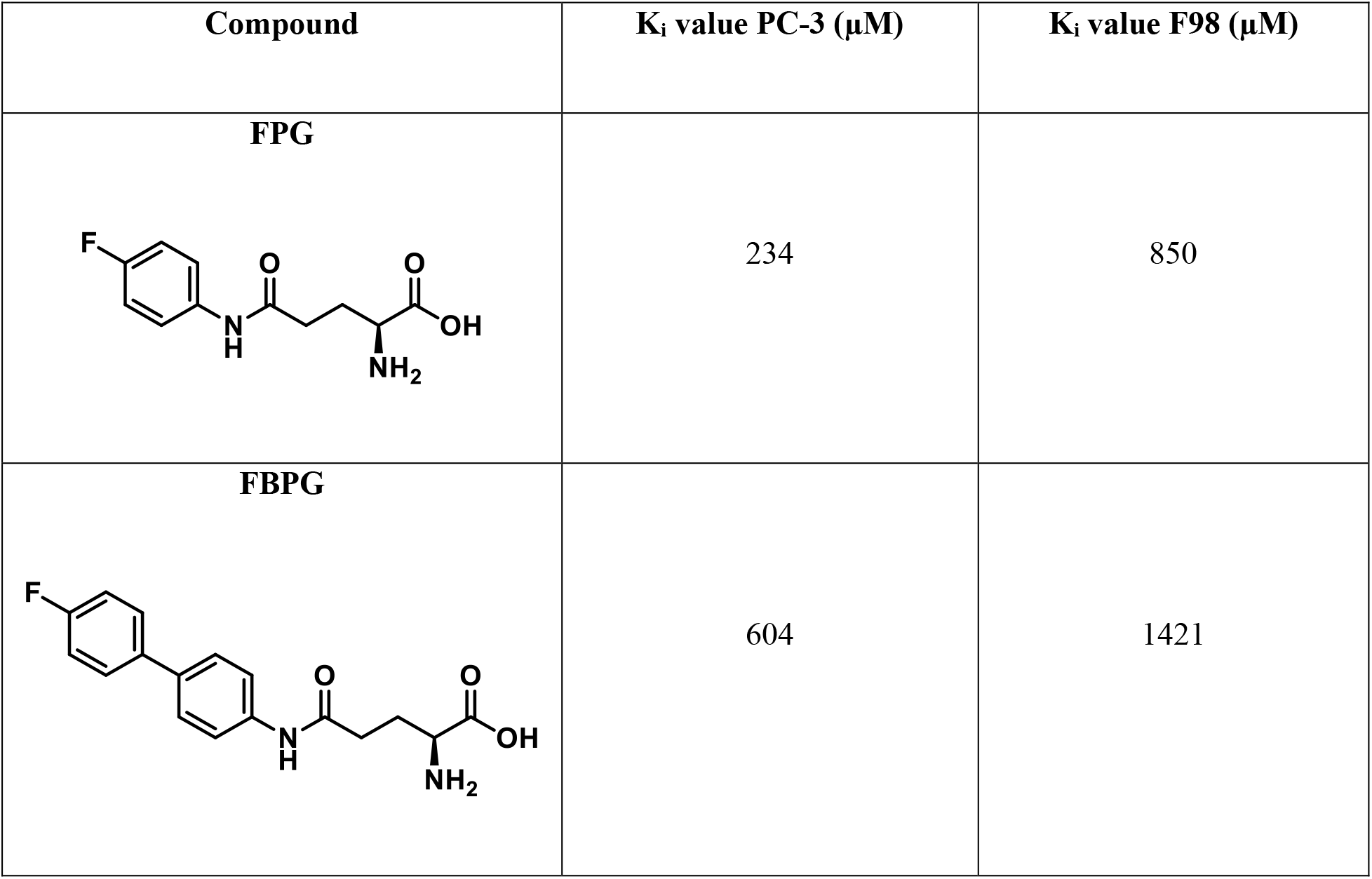
K_i_ values of FPG and FBPG for the ASCT-2 transporter on both the PC-3 and F98 cell line. Km (values in absence of cold product) and K_m,app_ (apparent values in presence of cold product) data were used to calculate the K_i_ values according to the formula: *Ki* = [*I*]/((*Km_app_*/*Km*) –1) with [I] being the concentration of inhibitor.

| Compound | $K_i$ value PC-3 (μM) | $K_i$ value F98 (μM) |
| --- | --- | --- |
| **FPG** | 234 | 850 |
| **FBPG** | 604 | 1421 |

#### Inhibition [^3^H]-l-glutamine uptake in knock-out cell line

Table 2^2^ compares the percentual inhibition of [^3^H]glutamine uptake between the ASCT-2 expressing cell line F98 and the HEK 293 knock-out cell line that does not express ASCT-2. Glutamine is transported across the cell membrane by multiple transporters, after knock-out of ASCT-2 in the HEK 293 cell line, the other transporter families still preserve their function. Both compounds inhibit the [^3^H]glutamine transport in the absence of the ASCT-2 transporter. As such, they inhibit the other glutamine transporting proteins, indicating that they are not specific for ASCT-2.

**Table 2:** Inhibition of [^3^H]glutamine uptake by FPG and FBPG in the F98 cell line and the ASCT-2 knockout cell line.

| Cell line | FPG | FBPG |
| --- | --- | --- |
| F98 | $6 \pm 11\%$ | $13 \pm 11\%$ |
| HEK 293 KO | $40 \pm 7\%$ | $42 \pm 9\%$ |

#### In vitro stability

Both [^18^F]FPG and [^18^F]FBPG remained stable in their formulation for at least 3 hours. No degradation products were found by means of analytical HPLC.

### d. In vivo experiments

#### PC-3 xenograft model

Semi-quantitative values for both radiotracers can be found in Table 3^3^. [^18^F]FPG shows moderate tumor uptake in the mice bearing PC-3 xenografts (see Figure 3). Only a very low uptake in the PC-3 xenograft was seen for [^18^F]FBPG (see Figure 3).

**Figure 3:**
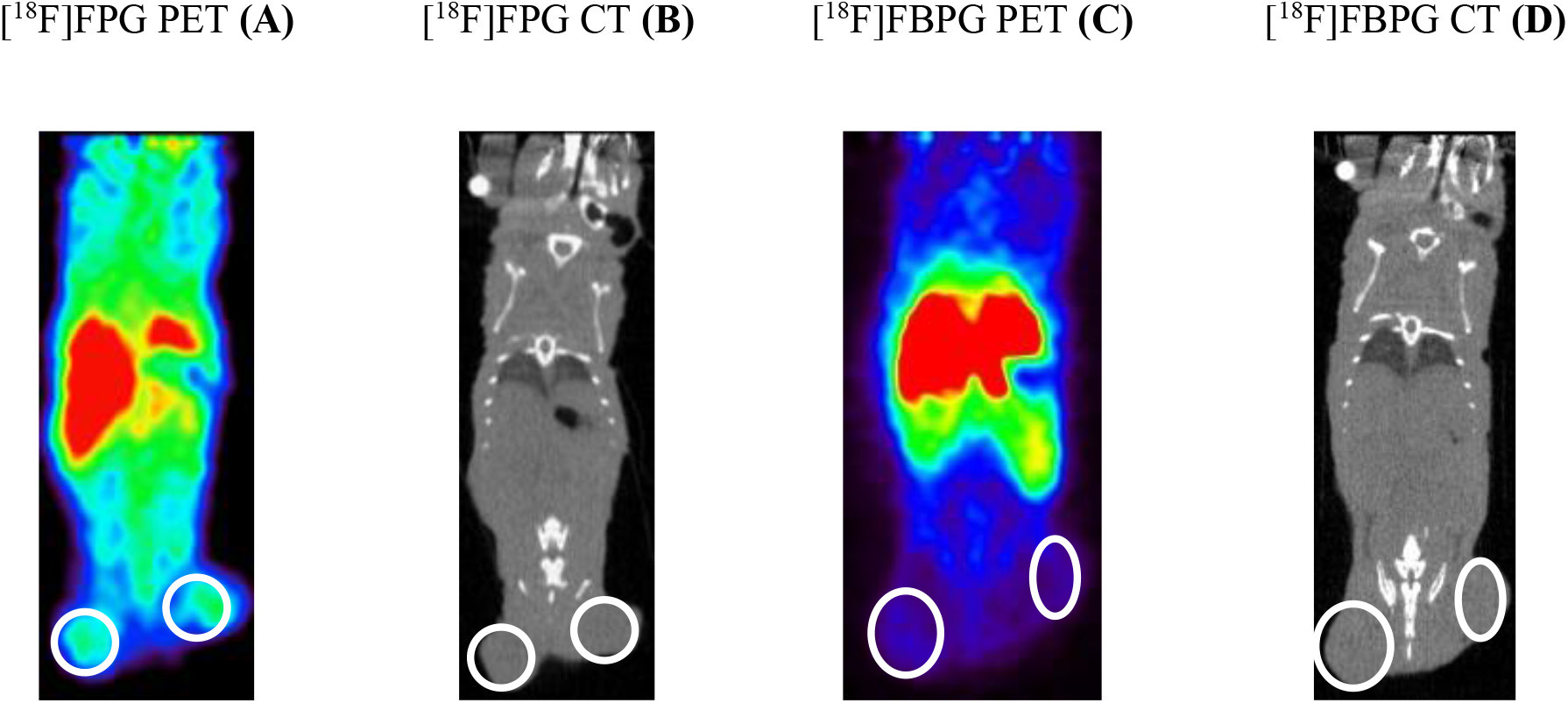
μPET images (**A** and **C**) and CT images (**B** and **D**) of the PC-3 bearing mouse xenografts, tumors are delineated with a white line. [^18^F]FPG has a moderate uptake in the tumors in both flanks (**A**), [^18^F]FBPG has only limited uptake (**C**) in both tumors.

**Table 3:** Semi-quantitative analysis of [^18^F]FPG and [^18^F]FBPG uptake in the PC-3 xenograft. For both radiotracers, the mean tumor-to-muscle and mean standardized uptake value were calculated.

|  | FPG | FBPG |
| --- | --- | --- |
| $\text{T/M}_{\text{mean}}$ | $1.68 \pm 0.24$ | $1.20 \pm 0.15$ |
| $\text{SUV}_{\text{mean}}$ | $0.53 \pm 0.25$ | $0.17 \pm 0.04$ |

#### Orthotopic F98 model

Glioblastoma growth in the right frontal lobe was confirmed on both T2-weighted and contrast-enhanced T1-weighted MRI, two weeks post-inoculation of F98 cells. Although both radiotracers enter the rat brain, no accumulation in the tumor was seen (see Figure 4).

**Figure 4:**
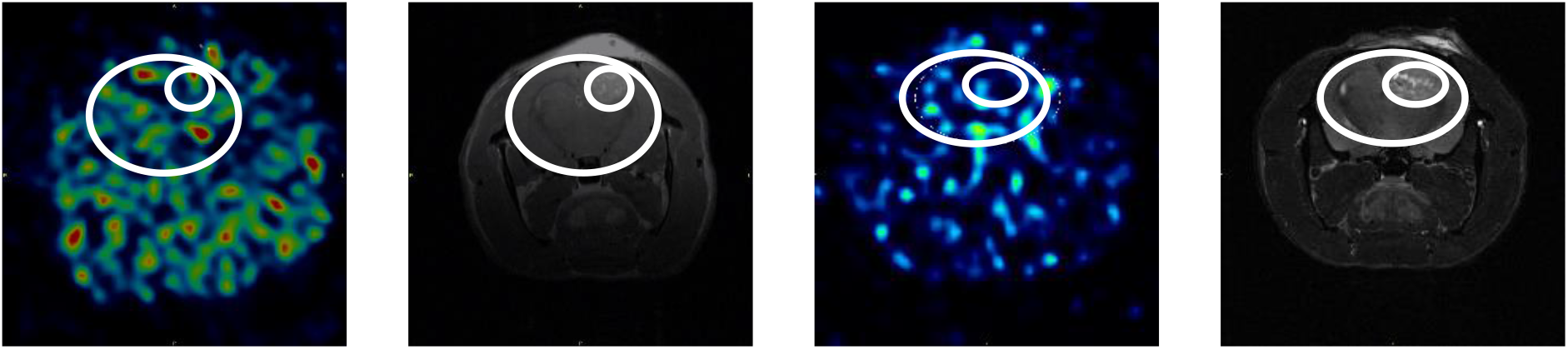
μPET images (**A** and **C**) and contrast-enhanced T1-weighted MR images (**B** and **D**) of F98 bearing rat brain (outer white circle). Macroscopical tumor growth (delineated by the inner white circle) was confirmed on the MRI images. Although both radiotracers enter the rat brain, no increased uptake in the tumor tissue was observed.

**Figure 5:**
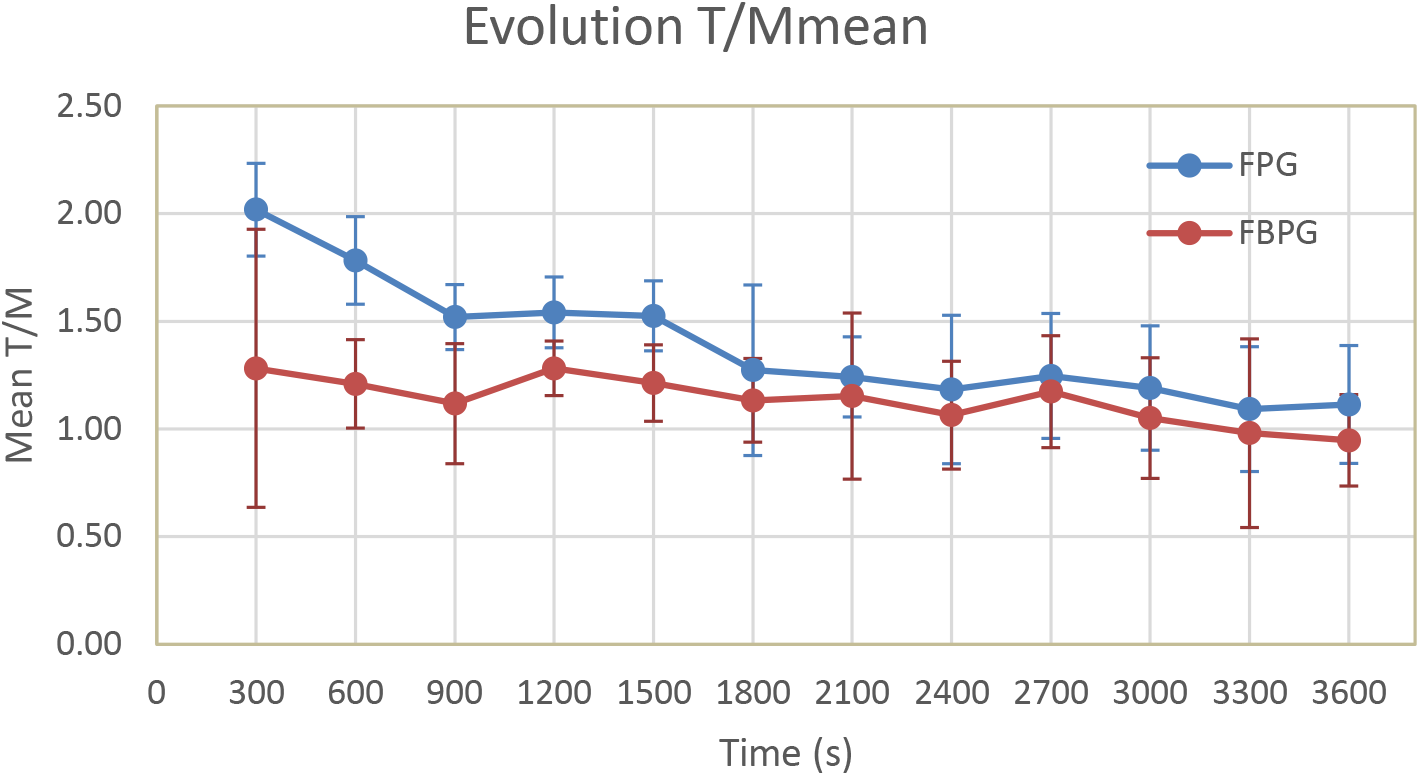
Evolution of the mean tumor-to-muscle ratio in function of the time for [^18^F]FPG (blue) and [^18^F]FBPG (red) in the PC-3 animal model.

#### Semi-quantitative analysis

The acquired PET data of [^18^F]FPG and [^18^F]FBPG were analyzed using the AMIDE and PMOD software. Semi-quantitative values, tumor-to-muscle (T/M) ratio and standard uptake value (SUV), were calculated and are depicted in Table 3. The T/M ratio for [^18^F]FPG peaks at the beginning of the acquisition and declines over time. [^18^F]FBPG has a tumor-to-muscle ratio close to one which does not change significantly over time.

## 4. Discussion

The metabolic importance of glutamine in tumor biology raised the overall interest for amino acid derivatives as molecular probes. Research has led to the development of both therapeutic and diagnostic compounds targeting their transporters and metabolic enzymes^25–28^. Due to the drawbacks of [^18^F]FDG and the “glutamine addiction” of several tumors, different PET radiotracers based on the glutamine backbone have been developed^5–8,10–13,16,17^. In this study, we developed and characterized [^18^F]fluorophenylglutamine ([^18^F]FPG) and [^18^F]fluorobiphenylglutamine ([^18^F]FBPG), two novel fluorine-18 labeled glutamine derivatives. Synthesis of the reference products and the precursors were successful. A satisfying yield (± 75%) was obtained for the synthesis of the cold reference products while only a moderate yield (± 58%) was found for the precursor synthesis. The introduction of unprotected hydroxyl groups, necessary for radiolabeling, results in a reduction of the synthesis yield. Masking the hydroxyl group (with a *tert*-butyldimethylsilyl-group) could possibly increase the yields of the amide formation. Deprotection of the amine and carboxylic acid with aqueous HCl led to impurities. We hypothesize that hydrolysis of the anilide is responsible for this degradation^29^. Removing the protecting groups with 4M HCl in dioxane resulted in pure products, that could be used without any further purification.

To obtain the fluorinated glutamine based PET tracers we firstly attempted to displace a nitro-group with fluoride-18 for the development of [^18^F]FBPG. However, due to the electrochemical properties of the compound the direct aromatic labelling was not feasible. Direct fluorination of the nitro moiety is possible, but requires harsh labelling conditions^30^. In addition, the electrochemical properties are very important for direct aromatic labelling, this can be seen in the dramatic changes of labelling yields of [^18^F]MPPF depending on the position of the electron withdrawing groups^31^. In neither [^18^F]FPG nor [^18^F]FBPG, electron withdrawing groups are present. Moreover, the amide and phenyl group were orientated *para* relative to the fluorine function as electron donating groups, further complicating the possibility for direct aromatic fluorination. Therefore, labelling the hydroxy functionalized precursor with the aid of the ruthenium complex was attempted. The ruthenium complex withdraws enough electron density from the phenol group through *η*^6^ *π*-coordination enabling labelling with fluoride-18 *via* a Meisenheimer complex even in the presence of electron donating groups.

LogD_7.4_ values of −0.70 and 1.11 were measured for [^18^F]FPG and [^18^F]FBPG, respectively. These values are much higher than the −3.64 logD_7.4_ value reported for glutamine^32^. This is explained by the introduction of the lipophilic side-chain to the glutamine. The implementation of a fluorophenyl group increased the lipophilicity with a hundredfold for [^18^F]FPG, the addition of a second phenylgroup in [^18^F]FBPG resulted in another thirtyfold increase. The concentration dependency experiments showed differences in affinity of both tracers towards the human PC-3 cells (Ki FPG= 234 μM; Ki FBPG= 604 μM) and rat F98 cells (Ki FPG= 850 μM; K_i_ FBPG= 1421 μM). This suggests conformational differences between rat and human ASCT-2 at the binding domain. This is supported by the low identity degree (14% as opposed to the 79% sequence identity) of both transporters and the previously reported difference in binding towards rat and human ASCT-2 of the 2-amino-4-bis(aryloxybenzyl)aminobutanoic acids (>50-fold difference)^33,34^. The difference in binding towards the ASCT-2 receptor in rat and human cells was confirmed by the *in vivo* experiments. With [^18^F]FPG, both tumors in the PC-3 mouse model were clearly visualized, whereas no tumor uptake was seen on PET images in the F98 rat model. In contrast to [^18^F]FPG, the discrepancy is not that clear with [^18^F]FBPG, where tumor delineation is not feasible in both tumor xenografts.

Because the μPET images with [^18^F]FPG provide a clear visualization of the tumor, we believe that this tracer is a good lead for the development of a radiotracer with high uptake in tumor tissue *in vivo*. Further increasing the affinity of the imaging probe towards ASCT-2 would be beneficial for visualizing malignancies. Work by Esslinger et al. emphasized the influence of arene substitutions on the binding properties^21^. Exploring a range of substitutions on the aromatic ring could provide more insight into the exact configuration of the lipophilic pocket described by several authors^21–23^. In this work, we developed a biphenyl analogue to exploit the lipophilic pocket and thereby increasing the affinity. However, the introduction of the second aromatic ring had a detrimental effect on the binding towards the ASCT-2 transporter. Therefore, we believe that the lipophilic pocket is limited in size and by introducing a second ring, the probe gets pushed out of the binding site, explaining the loss of affinity. Future work should focus on exploring the substitutions of the arene group. As such, increased affinities can be obtained which should result in higher *in vivo* uptake in the tumor tissue.

## 5. Conclusion

We investigated the imaging potential of two novel PET radiotracers [^18^F]FPG and [^18^F]FBPG. The direct aromatic fluorination with fluoride-18 was succesful using a ruthenium-based method, subsequent deprotection leads to the desired radiotracers. *In vitro* inhibition experiments showed a moderate and low affinity of fluorophenylglutamine and fluorobiphenylglutamine, respectively, towards the human ASCT-2 transporter. Both compounds had a low affinity towards the rat ASCT-2 transporter. These results were endorsed by the *in vivo* experiments with low uptake of both tracers in the F98 rat xenograft, low uptake of [^18^F]FBPG in the mice PC-3 xenograft and a moderate uptake of [^18^F]FPG in the PC-3 tumors. We hypothesize that increasing the affinity of [^18^F]FPG by optimizing the substituents of the arene ring can result in a high-quality glutamine-based PET radiotracer.

## Acknowledgement

MHB gratefully acknowledges the financial support through the startup funds from the University of Arkansas and the NIH-NIGMS (GM132906).

## Supporting Information

Figure S1: Analytical HPLC run from [^19^F]FPG.

Figure S2: Analytical HPLC run from [^18^F]FPG.

Figure S3: Analytical HPLC run from [^19^F]FBPG.

Figure S4: Analytical HPLC run from [^18^F]FBPG.

## ABBREVIATIONS

ASCT-2: Alanine-serine-cysteine transporter-2
CT: Computed tomography
FDA: Federal drug agency
FDG: Fluorodeoxyglucose
FDOPA: 3,4-dihydroxy-6-fluoro-L-phenylalanine
FET: O-(2-fluoroethyl)-L-tyrosine
FPG: Fluorophenylglutamine
FBPG: Fluorobiphenylglutamine
GPNA: L-γ-glutamyl-p-nitroanilide
LAT-1: L-type amino acid transporter
MET: (S-methyl)-L-methionine
MRI: Magnetic resonance imaging
PET: Positron emission tomography
SUV: Standardized uptake value
T/M: Tumor-to-muscle

## List of Figure captions

Figure 6: FACS analysis of ASCT-2 presence on PC-3 and F98 cells. The top left figure represents surface staining (negative control). The top right figure is intracellular staining of PC-3 cells. The bottom left figure is surface staining of F98 cells (negative control). The bottom right figure is intracellular staining of F98 cells.

Figure 7: Concentration dependent uptake of [^3^H]glutamine in F98 cell line (top row) and PC-3 cell line (bottom row). Top left and bottom left graphs represent baseline uptake of [^3^H]glutamine. The middle graphs represent uptake in presence of FPG. Top and bottom right graphs show the uptake of [^3^H]glutamine in the presence of FBPG.

Figure 8: μPET images (**A** and **C**) and CT images (**B** and **D**) of the PC-3 bearing mouse xenografts, tumors are delineated with a white line. [^18^F]FPG has a moderate uptake in the tumors in both flanks (**A**), [^18^F]FBPG has only limited uptake (**C**) in both tumors.

Figure 9: μPET images (**A** and **C**) and contrast-enhanced T1-weighted MR images (**B** and **D**) of F98 bearing rat brain (outer white circle). Macroscopical tumor growth (delineated by the inner white circle) was confirmed on the MRI images. Although both radiotracers enter the rat brain, no increased uptake in the tumor tissue was observed.

Figure 10: Evolution of the mean tumor-to-muscle ratio in function of the time for [^18^F]FPG (blue) and [^18^F]FBPG (red) in the PC-3 animal model.

## Footnotes

1 See last page

2 See last page

3 See last page

