## Supporting Info for "Radiosynthesis, *in vitro* and preliminary *in vivo* evaluation of the novel glutamine derived PET tracers [^18^F]fluorophenylglutamine and [^18^F]fluorobiphenylglutamine"

**Contents:**

1. Figure S1: Analytical HPLC run from [$^{19}$F]FPG.
2. Figure S2: Analytical HPLC run from [$^{18}$F]FPG.
3. Figure S3: Analytical HPLC run from [$^{19}$F]FBPG.
4. Figure S4: Analytical HPLC run from [$^{18}$F]FBPG

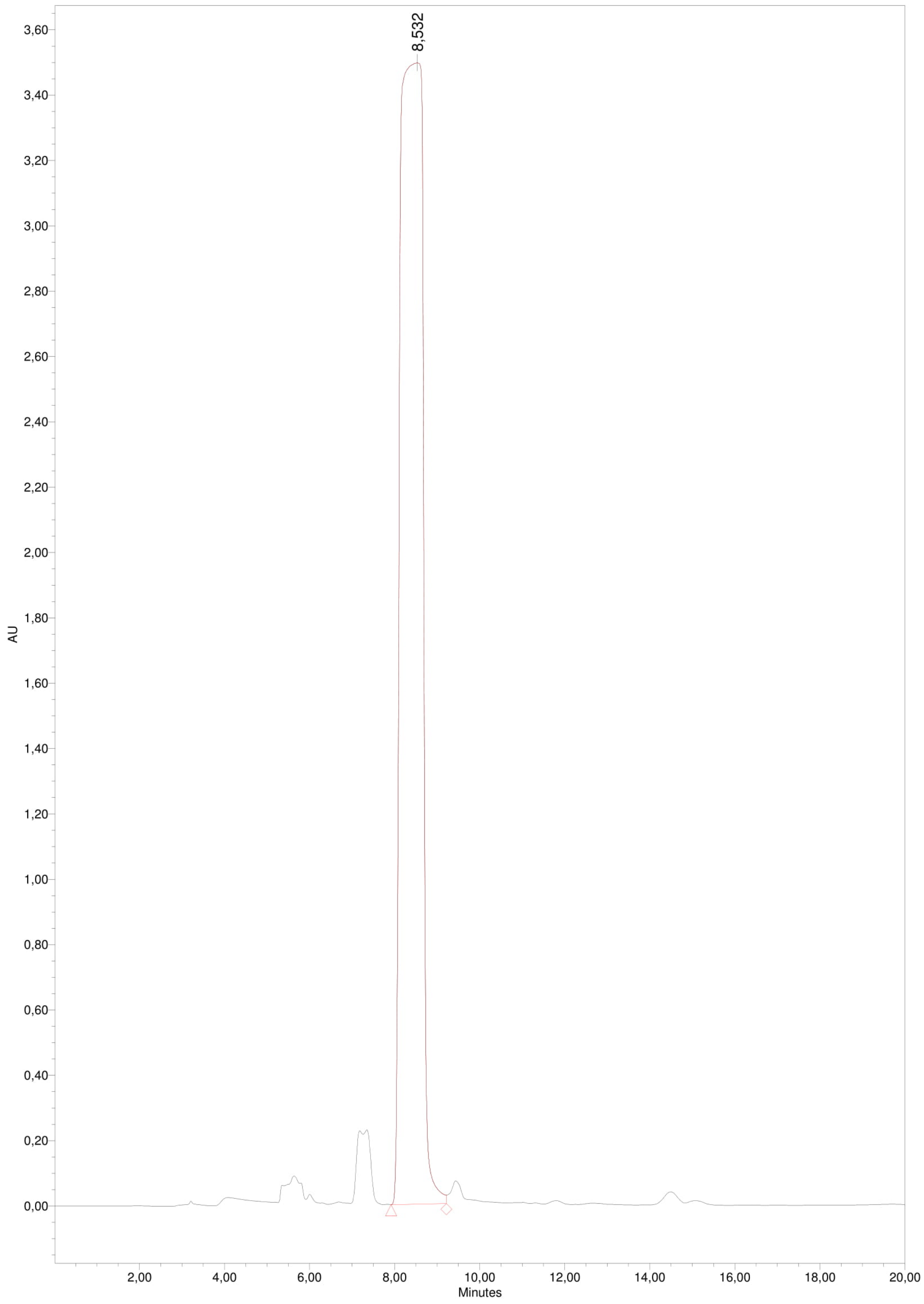


Figure S5: Analytical HPLC run from [$^{19}$F]FPG.

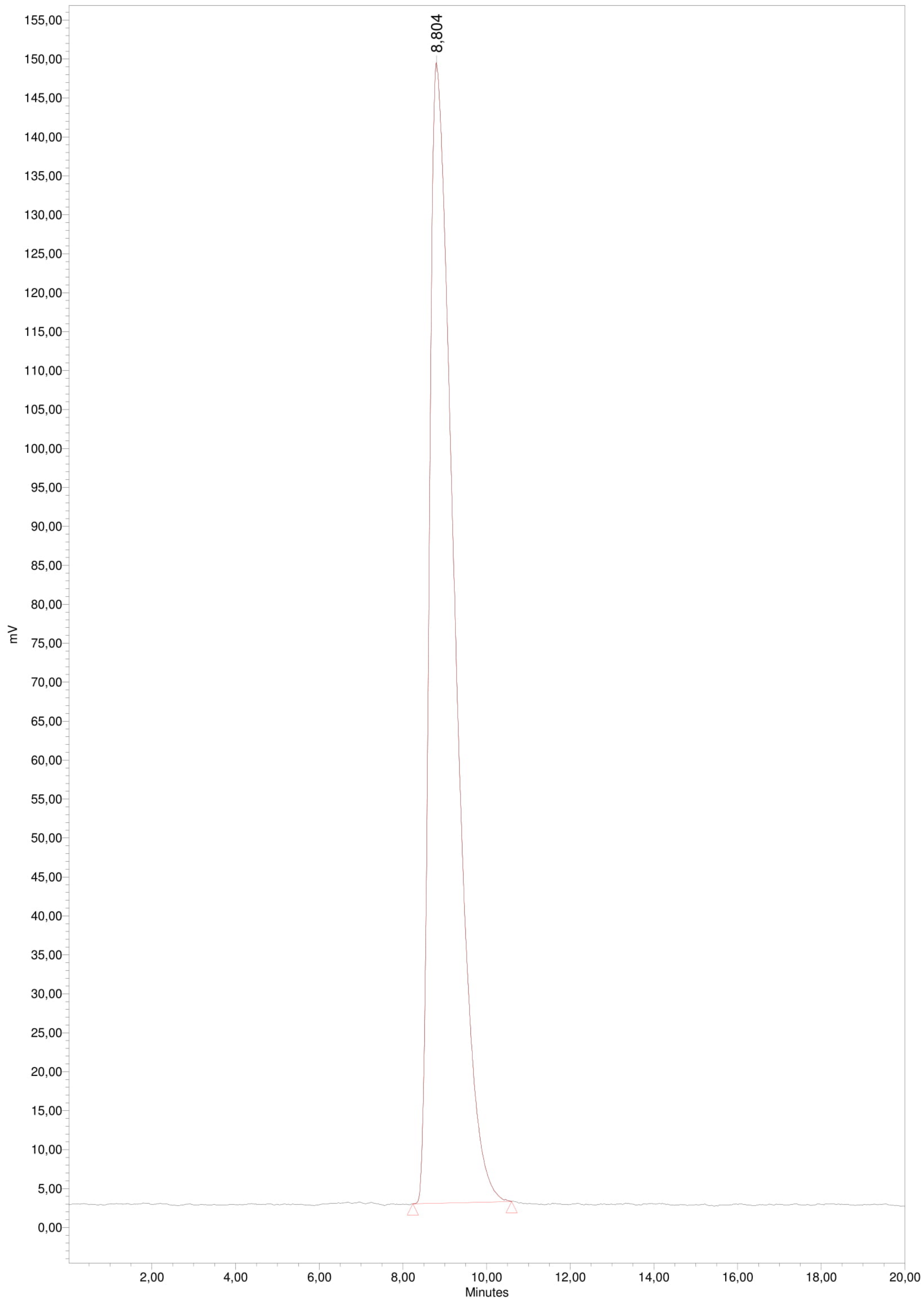


Figure S6: Analytical HPLC run from [$^{18}$F]FPG.

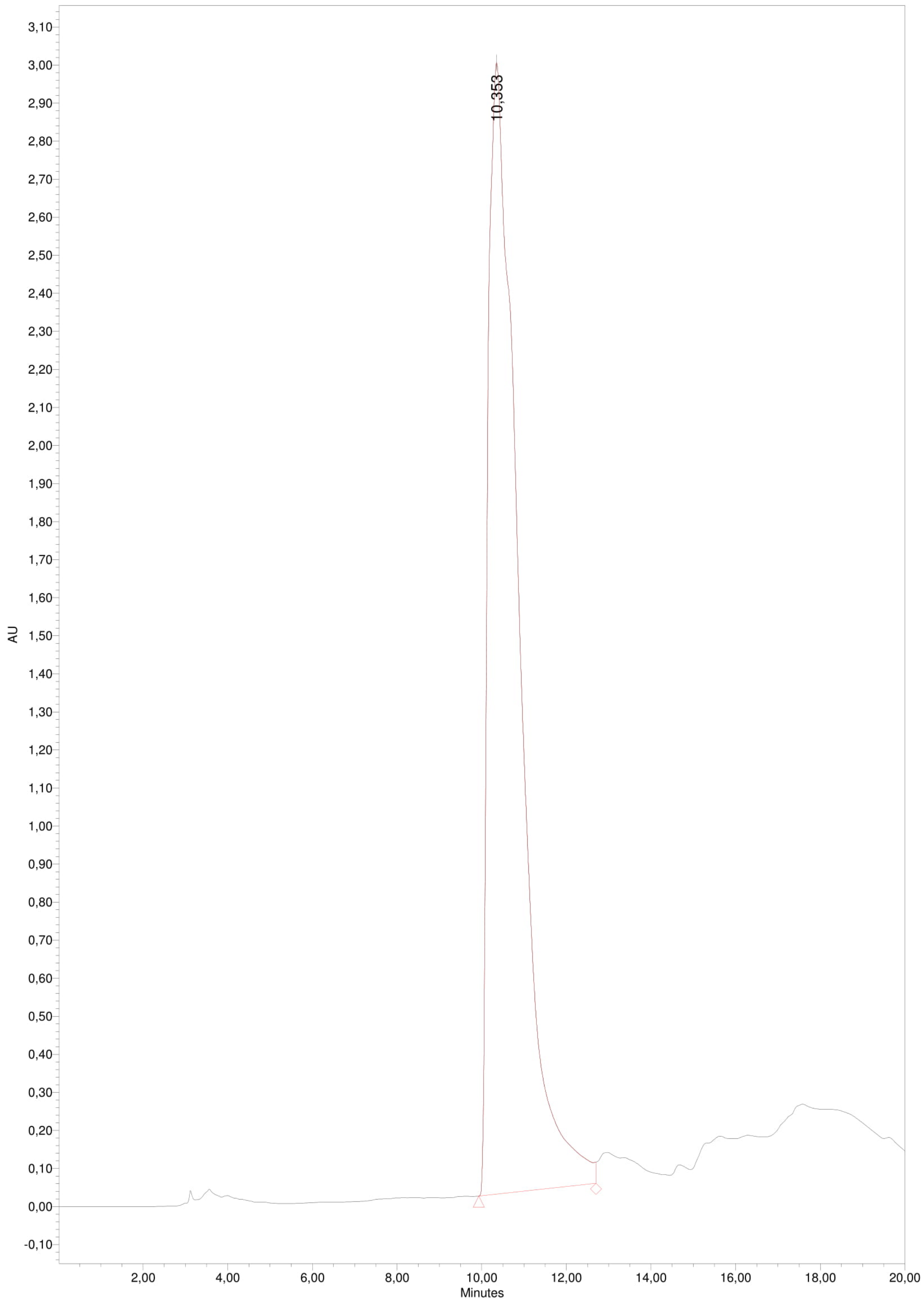


Figure S7: Analytical HPLC run from [$^{19}$F]FBPG.

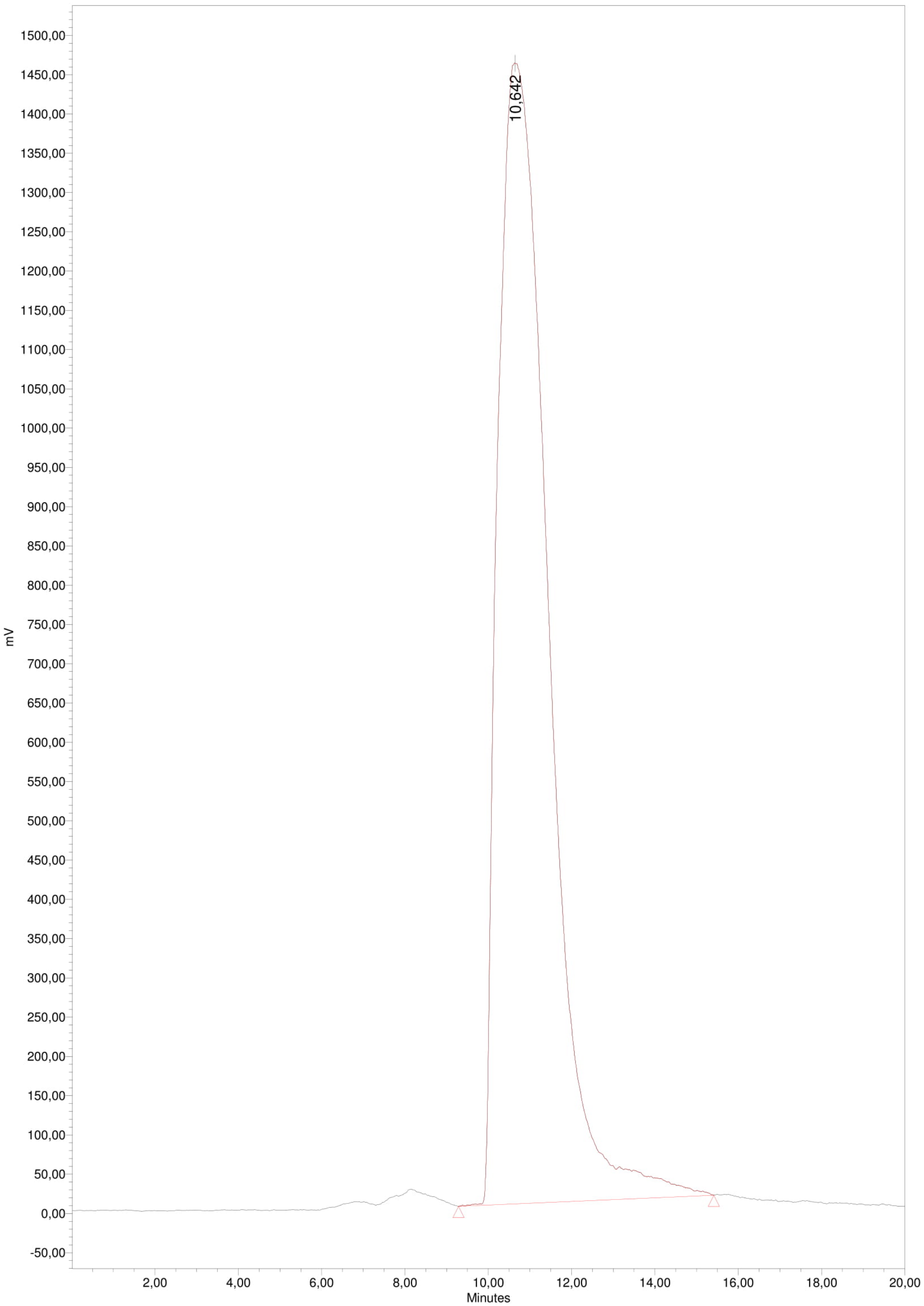


Figure S8: Analytical HPLC run from [$^{18}$F]FBPG
